# Rapoport’s rule in the marine realm: wrong axis, right pattern

**DOI:** 10.64898/2025.12.30.696717

**Authors:** Gabriel Reygondeau, Yulia Egorova

## Abstract

**Aim:** Rapoport’s rule posits that species’ range size increases with distance from benign conditions along environmental gradients. We asked whether, and along which axis, marine species obey Rapoport’s rule when ranges are quantified in three dimensions.

**Location:** Global ocean.

**Taxon:** > 20,000 marine species spanning pelagic and benthic habitats across major animal phyla and fishes.

**Methods:** We combined AquaMaps 2.0 / AquaX modelled distributions with independently curated depth limits from FishBase and SeaLifeBase to estimate latitudinal and vertical (bathymetric) ranges for each species. We quantified latitudinal range versus absolute mid-latitude and depth range versus mid-depth, and repeated analyses by habitat and taxonomic group. We used linear and polynomial regressions and Gaussian mixture models in range–gradient space to identify and compare alternative Rapoport regimes.

**Results:** Marine biodiversity exhibits a classic latitudinal diversity gradient and strong concentration of richness in the upper ocean, with broader ranges at higher latitudes and greater depths. Support for Rapoport’s rule is weak and inconsistent for latitudinal ranges, but strong and pervasive with depth, with correlations between vertical range and mid-depth frequently exceeding 0.8 across habitats and taxa. Latitudinal and vertical ranges are positively, but only moderately, coupled, with a small subset of “3-D generalists” spanning both large latitudinal and depth extents.

**Main conclusions:** The marine realm appears to obey Rapoport’s rule primarily along the vertical, rather than latitudinal, axis. Depth-structured environmental tolerance thus emerges as a dominant constraint on marine range limits and three-dimensional biodiversity gradients.

## Introduction

Rapoport’s rule links species’ range sizes to their position along geographical gradients: at higher latitudes, or at higher elevations on land, species tend to occupy broader geographic or elevational ranges than those closer to the equator or sea level (Gaston & Chown, 1999; G. C. Stevens, 1992). This pattern has been invoked to explain major features of terrestrial diversity, including the latitudinal diversity gradient and the dynamics of seasonal migration, under the idea that greater climatic variability at high latitudes selects for broader niche breadth and hence larger ranges (Pintor et al., 2015; Saupe et al., 2019). In the ocean, both the latitudinal diversity gradient and Rapoport’s rule have been tested, but support has been mixed. Several studies report weak or taxon-specific poleward increases in range size, and no single, robust marine analogue of the terrestrial pattern has emerged from analyses confined to latitude alone (Costello & Chaudhary, 2017; Fortes & Absalão, 2004; Kendall & Haedrich, 2006; G. Stevens, 1996).

Most of these studies, however, treat the ocean as a one-dimensional, horizontal system. They examine range sizes as a function of latitude, occasionally longitude, but rarely consider that the ocean is inherently three-dimensional and strongly structured in the vertical dimension (Longhurst, 2010). Depth underpins some of the steepest environmental gradients on the planet (from light and temperature to pressure, oxygen and resource supply) and these gradients are tightly linked to physiology, life history and dispersal (Álvarez-Noriega et al., 2020; Reygondeau et al., 2018). By analogy to the altitudinal expression of Rapoport’s rule in mountains, a “bathymetric Rapoport’s rule” in the sea is therefore conceptually plausible: species centered deeper in the water column might, in principle, occupy broader vertical ranges than shallow-water specialists, independently of any latitudinal trend (Costello et al., 2017). This raises a broader question that has rarely been addressed explicitly (Fortes & Absalão, 2010; Smith & Gaines, 2003): how does Rapoport’s rule behave in three-dimensional space (latitude and depth), and how are marine range sizes coupled to biodiversity patterns along these intersecting gradients?

We combine AquaMaps 2.0 (Reygondeau et al., 2025) outputs for over 20,000 marine species to estimate their latitudinal ranges, and pair these with independently curated minimum and maximum depth information from FishBase and SeaLifeBase to derive corresponding vertical (bathymetric) ranges. This joint dataset allows us to quantify latitudinal and vertical range sizes, and to relate them both to the central latitude and depth of species and to spatial patterns of species richness. In this letter, we first revisit the latitudinal form of Rapoport’s rule for marine species and then extend the analysis into the vertical dimension, testing whether Rapoport-like patterns are more clearly expressed with depth than with latitude, and how three-dimensional range structure aligns with the biodiversity gradient (species richness) of the global ocean.

## Material and Methods

### Data sources and range metrics

We compiled species-level latitudinal and vertical range information for marine taxa using AquaMaps 2.0 distribution outputs (Reygondeau et al., 2025) and depth traits from FishBase and SeaLifeBase. For each species, we first extracted the northernmost and southernmost predicted limits of its distribution from AquaMaps 2.0 outputs, calculated its latitudinal range as the difference between these two limits, and computed its latitudinal centroid (also referred as mid-latitude) as their arithmetic mean over their current distribution. To investigate Rapoport’s rule in relation to distance from the equator, we used the absolute value of this mid-latitude as the predictor, thereby collapsing hemispheres while retaining information on tropical versus temperate–polar position.

Vertical ranges were obtained independently of AquaMaps 2.0 using published trait information. For each species, we identified the shallowest and deepest realised occurrence depths reported in FishBase and SeaLifeBase, defined vertical range size as the difference between these two limits, and calculated mid-depth as their arithmetic mean.

All analyses were limited to species for which both latitudinal limits and depth limits were available (n =21,958). Species were additionally classified by broad habitat type (e.g., pelagic vs. benthic/demersal, following FishBase and SeaLifeBase) and by major taxonomic or functional group.

### Compute species Richness and Range

To quantify species richness gradients and characterize typical range extents across latitude and depth, we binned species by their mid-latitude and mid-depth. For the horizontal dimension, we empirically divided the latitudinal dimension in 0.25° band and based on the distribution of each species (from AquaMaps 2.0), we calculated the species richness as the number of species that fell inside each bin (Figure 1a). We then summarized range structure by computing the mean, minimum and maximum latitude over all species within each bin, which define an average latitudinal range envelope at each latitude (Figure 1b). For the vertical dimension, species were allocated to fixed depth bins according to their depth range, and we computed depth-specific species richness as the number of species whose depth range fell within each bin (every 10m from 0 to 200m, every 50m from 200m to 1000m, every 100m from 1000m to maximal depth) (Figure 2a). For each depth bin, we similarly calculated the mean, shallow and deep limits to describe an average vertical range envelope through the water column (Figure 2b). All richness and range-bin summaries were generated separately for pelagic versus benthic/demersal species (and for major taxonomic/functional groups where relevant), enabling comparison of horizontal and vertical gradients among habitats.

**Figure 1.**
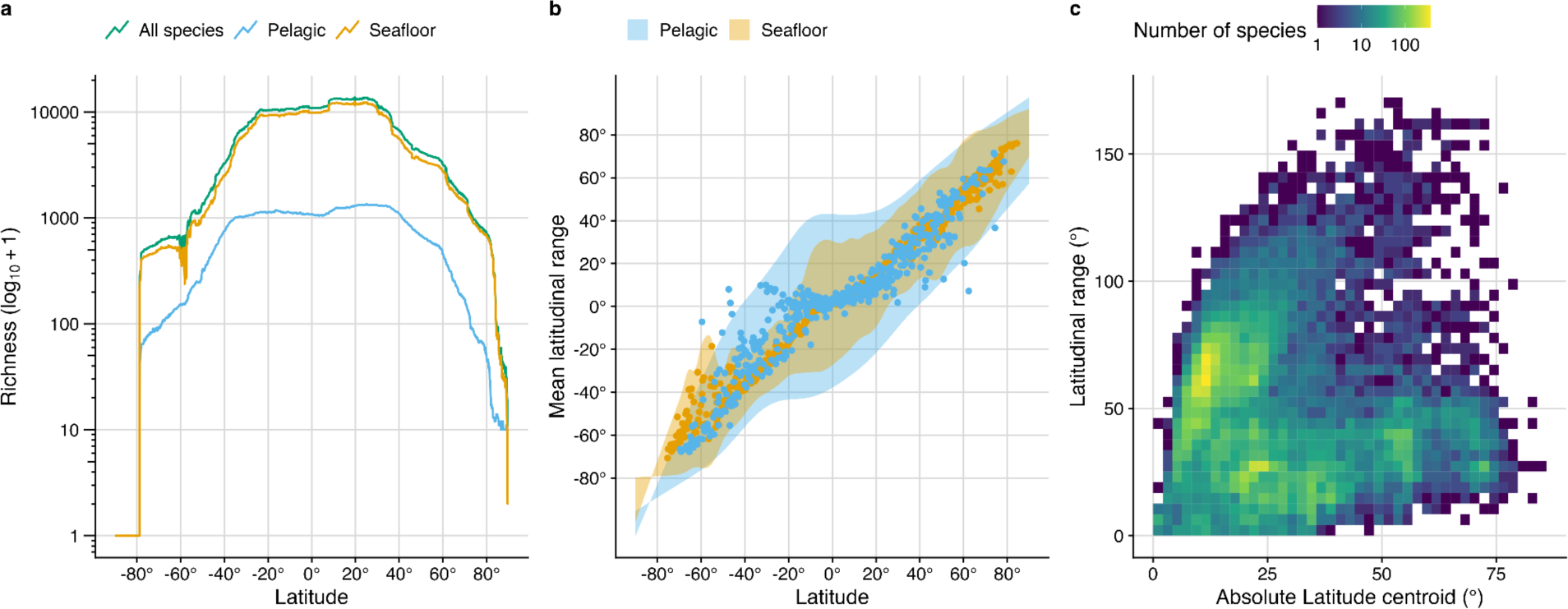
Latitudinal gradients in marine biodiversity and range size for all marine species, pelagic taxa and seafloor (benthic/demersal) taxa. (a) Latitudinal species richness (number of species per 0.25° band, log₁₀[richness + 1] on the y-axis) for all species (green), pelagic species (blue) and seafloor species (orange). (b) Mean latitudinal range (±1 SD ribbon) of pelagic (blue) and seafloor (orange) species as a function of latitude, showing the systematic increase in range size from equatorial to high latitudes. (c) Two-dimensional density of species’ latitudinal range size against the absolute latitude of the range centroid; colours indicate the number of species per bin (from purple = few to yellow = many), highlighting the concentration of many small-ranged species at low latitudes and a sparser cloud of large-ranged species extending into higher latitudes.

**Figure 2.**
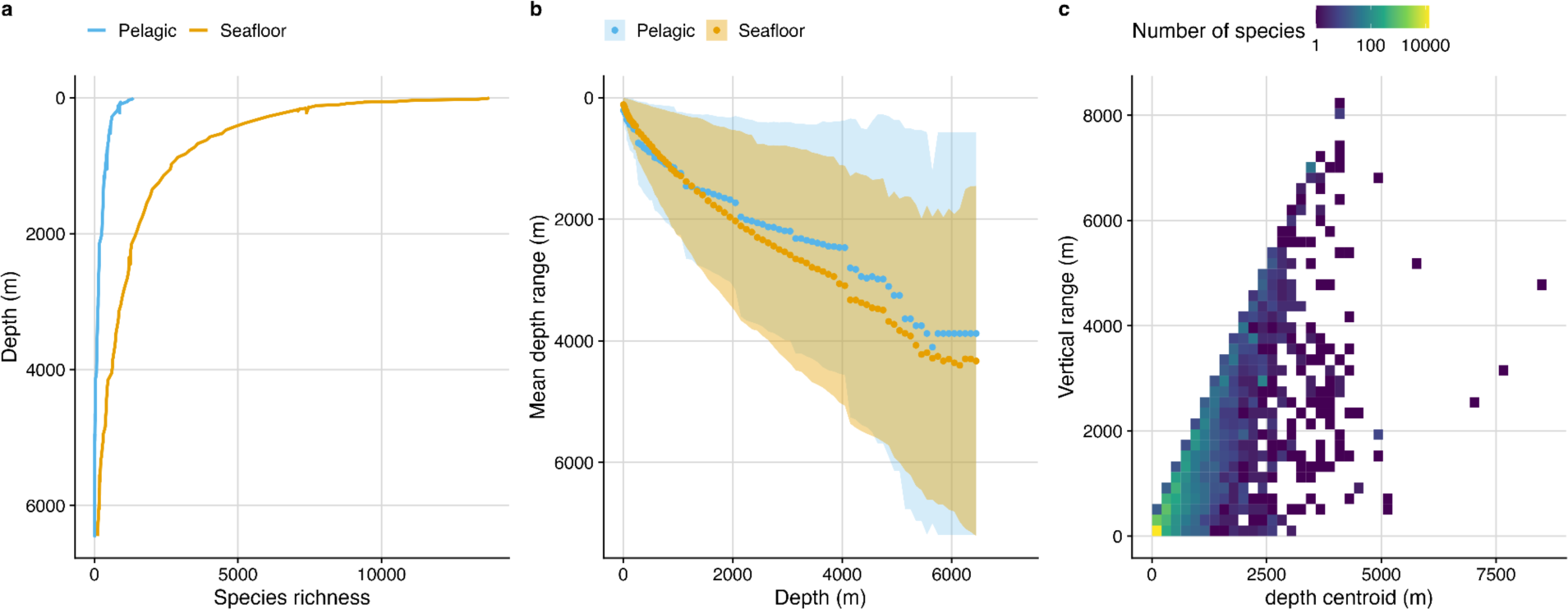
Vertical gradients in marine biodiversity and range size for pelagic and seafloor taxa. (a) Depth distribution of species richness (number of species per depth bin) for pelagic (blue) and seafloor (orange) species, showing the concentration of diversity in the upper few hundred metres and a rapid decline with depth. (b) Mean depth of species’ vertical ranges (points) and associated variability (±1 SD ribbons) as a function of bin mid-depth for pelagic (blue) and seafloor (orange) assemblages, illustrating systematic deepening and broadening of realised depth ranges. (c) Two-dimensional frequency distribution of species’ vertical range size versus the mid-depth of their range, with colours indicating the number of species per bin (from purple = few to yellow = many), highlighting that the widest vertical ranges occur at intermediate to bathyal depths rather than at the surface.

### Tests of Rapoport’s rule in horizontal and vertical dimensions

We tested Rapoport’s rule separately for the horizontal (latitudinal) and vertical dimensions. For the horizontal case, we related species’ latitudinal range size to their absolute mid-latitude, asking whether species occurring at higher absolute latitudes tend to have larger latitudinal ranges. For the vertical case, we related vertical range size to mid-depth, asking whether species centred deeper in the water column tend to have broader vertical ranges.

Analyses were conducted separately for each habitat type and taxonomic groups, to allow for distinct range–environment relationships among seafloor associated species versus pelagic and among different taxonomic/functional assemblages. For each subset, we fitted two simple linear models, where latitudinal or vertical range is the response variable and absolute mid-latitude or mid-depth is independent variable, respectively.

### Identifying multiple Rapoport regimes with Gaussian mixture models

Because Rapoport’s rule may not apply uniformly across all species, we explored whether distinct “regimes” of range–position relationships exist. To do this, we fitted two-component Gaussian mixture models (GMMs) (Ju & Liu, 2012) in the bivariate space of absolute mid-latitude and latitudinal range size. For each species, we modelled the joint distribution of species in absolute mid-latitude, latitudinal range space using an expectation–maximisation algorithm for a two-component GMM.

This approach partitions species into two latent clusters that differ in their combinations of geographic position and range size, allowing us to identify, for example, a set of species conforming more closely to a classical Rapoport pattern (larger ranges at higher absolute mid-latitude) versus another set with weaker or opposite trends. We then re-estimated range–position relationships within each Gaussian component to characterise these distinct Rapoport regimes.

### Three-dimensional range–richness space

To jointly examine horizontal and vertical dimensions of species ranges, we constructed a three-dimensional “range–range–richness” space defined by latitudinal range, vertical range, and local species richness. We binned species by latitudinal range and vertical range classes and counted the number of species within each two-dimensional bin. This yielded a three-dimensional surface where the horizontal axes represent latitudinal and vertical range sizes, and the vertical axis represents species richness (number of species).

## Results

Across the global ocean, species richness shows a clear latitudinal diversity gradient, with a broad peak in the tropics and subtropics and declining values toward both poles for all species, pelagic taxa, and seafloor (e.g., benthic/demersal) taxa (Fig. 1a). Richness is highest between about 10–30° in both hemispheres, and then drops steeply at higher latitudes, with the pelagic assemblage consistently poorer than the seafloor associated (benthic/demersal) assemblage at any given latitude. Mean latitudinal range size, in contrast, increases away from the equator for both habitats (Fig. 1b). Pelagic species tend to have slightly broader latitudinal ranges than seafloor species at mid–high latitudes, whereas both groups maintain a large pool of relatively small-ranged species in the tropics, as highlighted by the high-density cloud of short ranges at low latitude centroid in the 2D density plot (Fig. 1c). Together, these patterns are consistent with a “tropical cradle” of diversity composed of many small-ranged species, sitting atop a background of wider-ranging taxa that extend into temperate and polar waters.

Rapoport tests, however, show that this latitudinal cline in richness and the tendency for broader ranges at higher latitudes only weakly translate into a classic Rapoport relationship (Fig. 1c). Across all species, correlations between latitudinal range and absolute mid-latitude are small and slightly negative for both Pearson and Spearman metrics, and this pattern holds or strengthens for most major groups and for the benthic/demersal assemblage, whereas pelagic taxa show near-zero correlations (Table 1). In other words, although higher-latitude communities are dominated by broader-ranging species (highest range are found in the temperate area, Fig. 1b) and low latitudes concentrate many small-ranged taxa, the increase in range size with absolute latitude is neither strong nor universal enough to support a simple, monotonic Rapoport rule in the horizontal dimension. Instead, the latitudinal diversity gradient appears to emerge from a mixture of widespread taxa and multiple pools of small-ranged species whose distributions are structured by processes more complex than a single global range–latitude relationship.

**Table 1.**
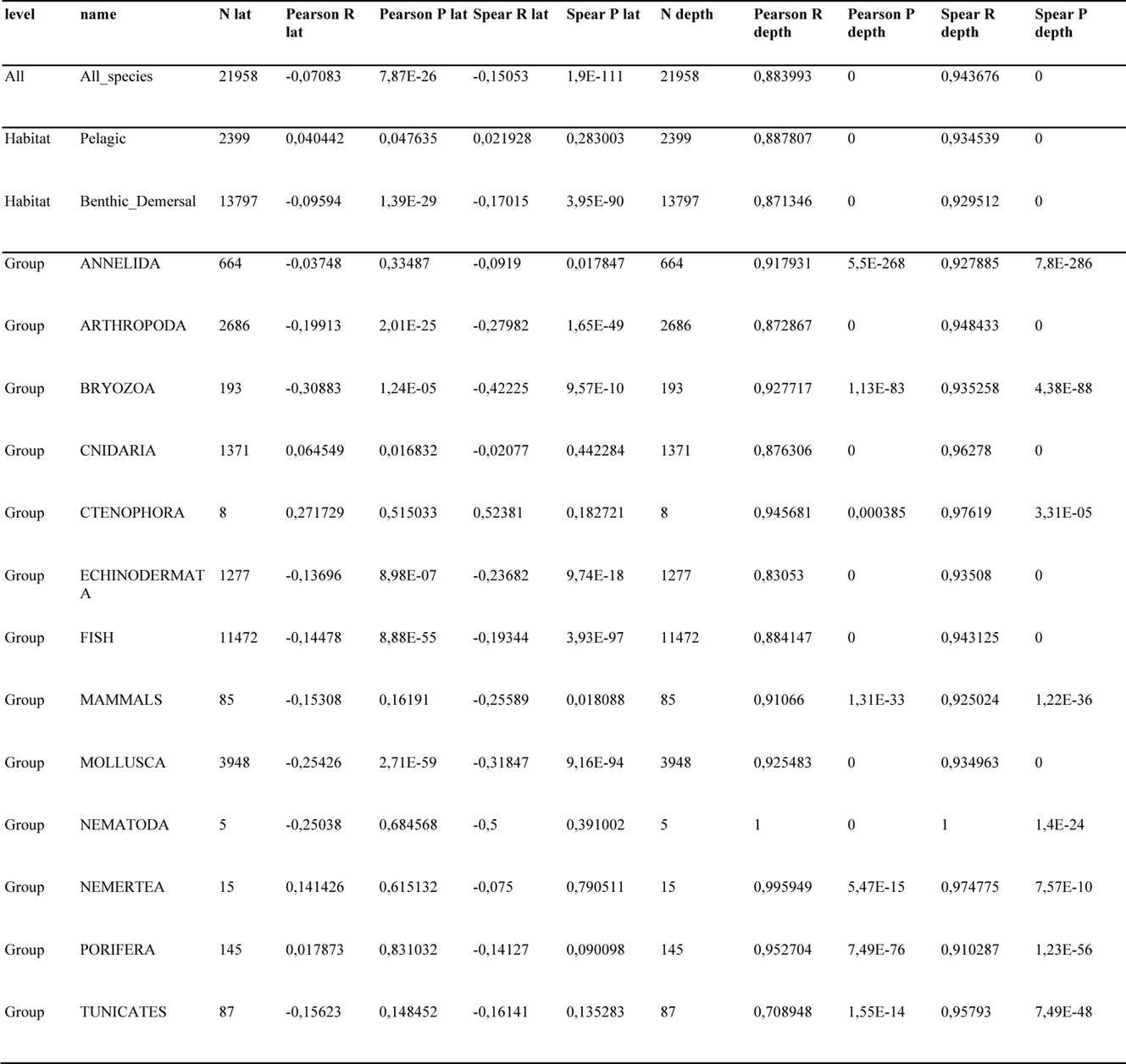
Summary of Rapoport’s rule tests for latitudinal and vertical range size across all marine species, habitat types and taxonomic groups. For each subset (all species, pelagic, benthic/demersal, and each taxonomic groups), we report the number of species (N), Pearson correlation coefficient (r) and Spearman rank correlation coefficient (ρ) between (i) latitudinal range and absolute mid-latitude, and (ii) vertical range and mid-depth, together with associated two-tailed P-values. Positive, significant coefficients indicate support for Rapoport’s rule in the corresponding dimension.

| level | name | N lat | Pearson R lat | Pearson P lat | Spear R lat | Spear P lat | N depth | Pearson R depth | Pearson P depth | Spear R depth | Spear P depth |
| --- | --- | --- | --- | --- | --- | --- | --- | --- | --- | --- | --- |
| All | All_species | 21958 | -0,07083 | 7,87E-26 | -0,15053 | 1,9E-111 | 21958 | 0,883993 | 0 | 0,943676 | 0 |
| Habitat | Pelagic | 2399 | 0,040442 | 0,047635 | 0,021928 | 0,283003 | 2399 | 0,887807 | 0 | 0,934539 | 0 |
| Habitat | Benthic_Demersal | 13797 | -0,09594 | 1,39E-29 | -0,17015 | 3,95E-90 | 13797 | 0,871346 | 0 | 0,929512 | 0 |
| Group | ANNELIDA | 664 | -0,03748 | 0,33487 | -0,0919 | 0,017847 | 664 | 0,917931 | 5,5E-268 | 0,927885 | 7,8E-286 |
| Group | ARTHROPODA | 2686 | -0,19913 | 2,01E-25 | -0,27982 | 1,65E-49 | 2686 | 0,872867 | 0 | 0,948433 | 0 |
| Group | BRYOZOA | 193 | -0,30883 | 1,24E-05 | -0,42225 | 9,57E-10 | 193 | 0,927717 | 1,13E-83 | 0,935258 | 4,38E-88 |
| Group | CNIDARIA | 1371 | 0,064549 | 0,016832 | -0,02077 | 0,442284 | 1371 | 0,876306 | 0 | 0,96278 | 0 |
| Group | CTENOPHORA | 8 | 0,271729 | 0,515033 | 0,52381 | 0,182721 | 8 | 0,945681 | 0,000385 | 0,97619 | 3,31E-05 |
| Group | ECHINODERMAT A | 1277 | -0,13696 | 8,98E-07 | -0,23682 | 9,74E-18 | 1277 | 0,83053 | 0 | 0,93508 | 0 |
| Group | FISH | 11472 | -0,14478 | 8,88E-55 | -0,19344 | 3,93E-97 | 11472 | 0,884147 | 0 | 0,943125 | 0 |
| Group | MAMMALS | 85 | -0,15308 | 0,16191 | -0,25589 | 0,018088 | 85 | 0,91066 | 1,31E-33 | 0,925024 | 1,22E-36 |
| Group | MOLLUSCA | 3948 | -0,25426 | 2,71E-59 | -0,31847 | 9,16E-94 | 3948 | 0,925483 | 0 | 0,934963 | 0 |
| Group | NEMATODA | 5 | -0,25038 | 0,684568 | -0,5 | 0,391002 | 5 | 1 | 0 | 1 | 1,4E-24 |
| Group | NEMERTEA | 15 | 0,141426 | 0,615132 | -0,075 | 0,790511 | 15 | 0,995949 | 5,47E-15 | 0,974775 | 7,57E-10 |
| Group | PORIFERA | 145 | 0,017873 | 0,831032 | -0,14127 | 0,090098 | 145 | 0,952704 | 7,49E-76 | 0,910287 | 1,23E-56 |
| Group | TUNICATES | 87 | -0,15623 | 0,148452 | -0,16141 | 0,135283 | 87 | 0,708948 | 1,55E-14 | 0,95793 | 7,49E-48 |

In addition, the bivariate density of range size against absolute latitudinal centroid (Fig. 1c) is clearly not single-mode but organised into two arms. A Gaussian mixture model with two components fitted in absolute mid-latitude and range space partitions the data into two similarly sized clusters (N = 11,400 and 10,600 species). In the first cluster (Annex 1), which is strongly dominated by tropical species (82% tropical, 15% temperate, 2% polar), latitudinal range increases steeply with absolute latitude (slope = 1.6° range per 1° latitude, Pearson r = 0.72). The second cluster contains a much larger fraction of extra-tropical species (68% temperate and polar) and exhibits a shallower but still positive relationship (slope = 0.4, Person r = 0.50). Ecologically, the “steep arm” (Annex 1, red cluster) corresponds mainly to tropical-centred taxa whose distributions extend into subtropical and temperate zones, whereas the “shallow arm” is enriched in temperate and polar benthic/demersal assemblages with already broad ranges that expand more slowly with latitude. The weak global Rapoport signal in Table 1 therefore reflects the superposition of these two overlapping regimes rather than a single universal range–latitude slope.

Depth-wise, marine biodiversity shows a steep, non-linear decline with increasing distance from the surface. Species richness for both pelagic and seafloor assemblages is concentrated in the upper few hundred metres, with pelagic richness peaking in the epipelagic and dropping to near zero by ∼1,000–1,500 m, whereas seafloor richness declines more gradually along the continental slope but is already low beyond ∼3,000 m (Fig. 2a). Mean vertical position and range breadth co-vary strongly with depth: for both habitats, the centroid of species’ depth ranges deepens from shelf to bathyal zones, and the average vertical extent of these ranges broadens from a few tens of metres in shallow waters to several hundred or even thousands of metres at intermediate and bathyal depths (Fig. 2b). The two-dimensional frequency distribution of vertical range versus mid-depth reveals a dense wedge of shallow species with narrow vertical ranges, and a sparser but pronounced tail of “deep generalists” that combine kilometre-scale depth ranges with mid-depths between ∼1,000 and 4,000 m (Fig. 2c).

Tests confirm that Rapoport’s rule is strongly expressed in the vertical dimension, in contrast to the weak and inconsistent latitudinal signal (Table 1). Across all species, correlations between vertical range and mid-depth are very strong and positive (Pearson r = 0.88; Spearman r = 0.94), and this pattern persists when analyses are split by habitat and by major taxonomic group (Table 1). Both pelagic and benthic/demersal species show similarly high vertical Rapoport correlations (Pearson r = 0.87–0.89; Spearman r = 0.93–0.94), and most groups with sufficient sample sizes (e.g. fishes, molluscs, arthropods, echinoderms, cnidarians) all display Pearson and Spearman r values > 0.8. By contrast, the corresponding tests for latitudinal range versus absolute mid-latitude yield weak, often slightly negative correlations for the full data set and for most groups, with pelagic species in particular showing near-zero relationships.

Across all species, latitudinal and vertical range sizes were positively but only moderately correlated (Pearson r = 0.24, Spearman r = 0.14, N = 22,000), consistent with the joint distribution in Figure 3 where a dense core of small-ranged species is accompanied by a diffuse upper-right tail of “3-D generalists” that are broad in both dimensions. This coupling is slightly stronger for pelagic assemblages (Pearson r = 0.27, Spearman r = 0.34, N = 2,400) than for benthic/demersal species (Pearson r = 0.21, Spearman r = 0.10, N = 13,800), suggesting that water-column taxa are more likely to expand horizontally as their depth envelopes broaden (Table 1). At the group level, the pattern is heterogeneous but broadly similar: most major clades show weak-to-moderate positive Pearson correlations between depth and latitudinal range (e.g. fish Pearson r = 0.26, cnidarians r = 0.30, arthropods r = 0.21, annelids r = 0.21), with stronger associations in some groups such as echinoderms (r = 0.42) and marine mammals (r = 0.36), and near-zero or noisy relationships only in a few small groups with limited sample size (e.g. bryozoans, ctenophores). Overall, these results indicate that the processes that generate large vertical ranges (particularly in pelagic and deep-water taxa) do tend to promote broader latitudinal extents, but the effect is partial and group-dependent, reinforcing the view that vertical and horizontal Rapoport patterns are linked yet not governed by a single, universal mechanism.

**Figure 3.**
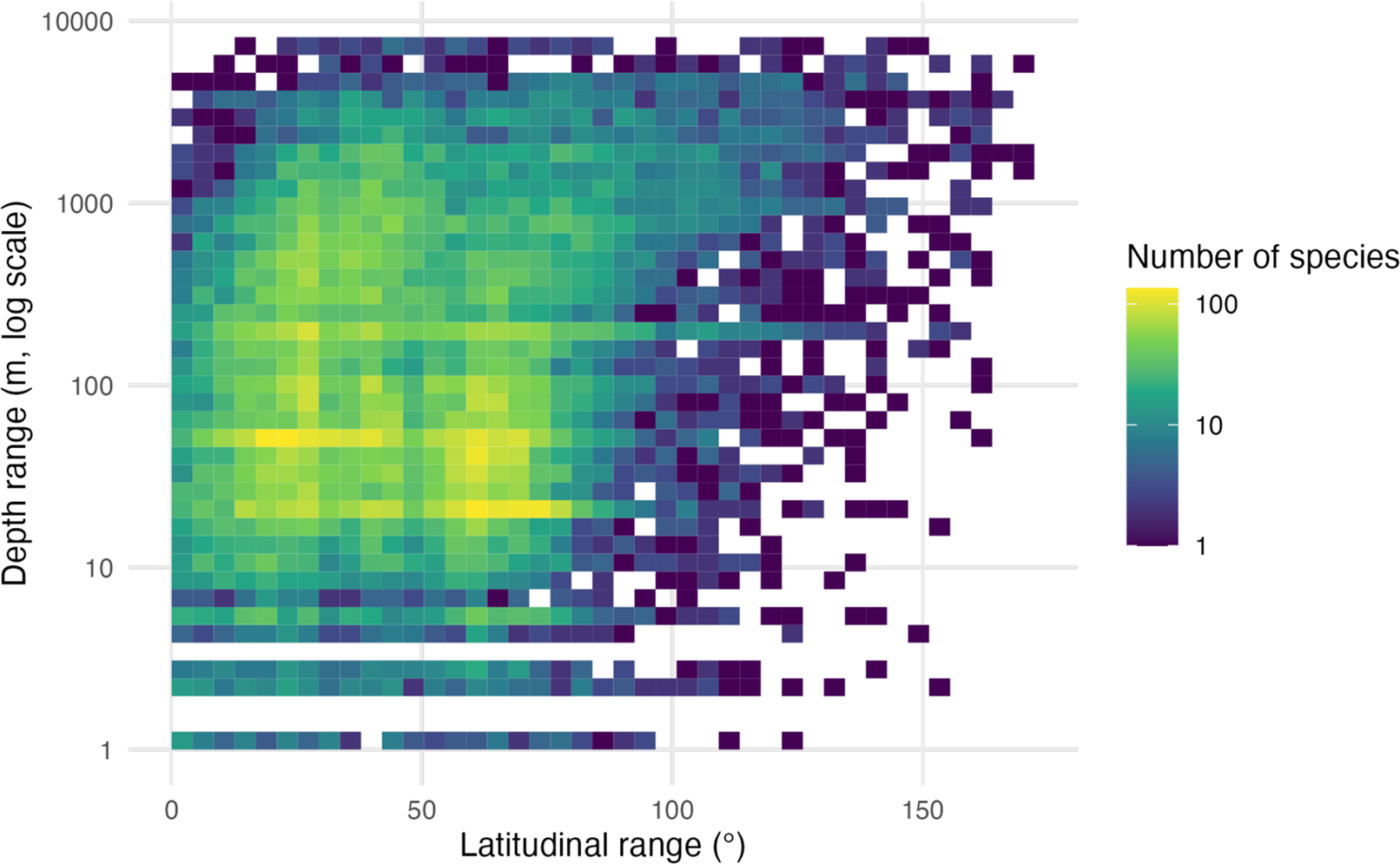
Coupling between latitudinal and vertical range size for marine species. Depth range (m, log₁₀ scale) is plotted against latitudinal range (°), with colours indicating the number of species in each two-dimensional bin (from dark = few to light = many). Most species occupy relatively small latitudinal and shallow depth ranges, whereas a sparse set of “3-D generalists” combines very wide depth ranges with broad latitudinal ranges, forming the long upper-right tail of the distribution.

## Discussion

Our analyses confirm that marine biodiversity exhibits a strong latitudinal diversity gradient and a pronounced concentration of richness in the upper ocean (Chaudhary et al., 2016), but they also show that range–environment relationships differ sharply between the horizontal and vertical dimensions (G. Stevens, 1996). Richness peaks at low to mid latitudes for both pelagic and seafloor assemblages and declines toward the poles, while vertical richness is tightly compressed into the epipelagic and upper slope, with only a modest “shoulder” at bathyal depths (Costello & Chaudhary, 2017; Owens & Rahbek, 2025). Superimposed on these gradients, range size varies systematically: latitudinal ranges tend to be broader away from the equator and vertical ranges broaden with increasing mid-depth. Yet, when tested explicitly, Rapoport’s rule is strongly supported only in the vertical dimension, and only weakly or inconsistently in the latitudinal dimension (Table 1).

The latitudinal results echo the mixed evidence reported from previous marine studies, which have generally found at best modest increases in range size with latitude, often restricted to particular clades or basins (Costello & Chaudhary, 2017; Fortes & Absalão, 2004, 2004; G. Stevens, 1996). At the scale of the global ocean, correlations between latitudinal range and absolute mid-latitude are close to zero or slightly negative, both for the full species pool and within most major taxonomic groups and habitats (Table 1). This is despite a clear macroecological pattern in which tropical and subtropical bands harbour large numbers of small-ranged species, while high-latitude assemblages are dominated by relatively broad-ranging taxa (Fig. 1b). Our mixture-model analysis helps reconcile these observations: rather than a single monotonic relationship, the cloud of points in range–latitude space decomposes into at least two regimes, one dominated by tropical species with steep increases in range size away from the equator, and another dominated by extra-tropical species whose ranges are already broad and expand more slowly with latitude (Fig. 1c). The superposition of these “arms” weakens any global, linear Rapoport signal.

By contrast, the vertical dimension conforms closely to the expectations of Rapoport’s rule. Across the combined data set and within most habitats and groups, correlations between vertical range and mid-depth are strong and positive, with slopes indicating that depth ranges increase by hundreds to thousands of metres from shelf to bathyal zones (Table 1). The two-dimensional frequency distributions show dense wedges of shallow, vertically narrow species and progressively more diffuse tails of deep, vertically broad “generalists”(Fig. 2b-c). This pattern holds for both pelagic and benthic/demersal species and remains robust when analyses are repeated within major clades.

Why should Rapoport’s rule emerge vertically but not horizontally in the sea? One possibility is that the processes constraining range limits along depth gradients are more strongly tied to environmental tolerance than those along latitude. Depth integrates a suite of environmental axes (light, pressure, temperature variability, oxygen, and, for demersal taxa, substrate type) that change predictably and largely monotonically with distance from the surface (Longhurst, 2010). Species that attain mid-depths in the mesopelagic or bathyal zones must tolerate a wide envelope of conditions if they are to complete their life cycles, which may naturally generate broader vertical ranges (Owens & Rahbek, 2025). By contrast, latitudinal gradients are intersected by strong regional heterogeneity in currents, productivity, boundary currents and basin geometry, and by the patchy distribution of continental margins (Chaudhary et al., 2016). These features can decouple thermal or seasonal regimes from simple distance to the equator and may allow small-ranged endemics to persist at many latitudes, especially in coastal and semi-enclosed seas.

Our analysis of coupling between vertical and horizontal range size offers further insight. Overall, depth and latitudinal ranges are positively but only moderately correlated, with stronger associations in pelagic assemblages and in some clades (e.g. echinoderms, mammals) than in others. This suggests that the mechanisms enabling broad vertical ranges (such as broad physiological tolerance, flexible life histories, or dispersive pelagic stages) do translate partially into broad latitudinal ranges, but only where circulation and habitat continuity permit. The existence of a small fraction of “3-D generalists” that combine kilometre-scale depth ranges with basin-scale latitudinal extents highlights this link (Fig. 3). However, the bulk of species remain restricted in at least one dimension, emphasising that the evolution of large ranges is not an automatic consequence of vertical tolerance and vice-versa.

These results have implications for how we interpret and model marine range shifts under climate change. If vertical range size is tightly linked to depth and is more strongly constrained by physiology and life history than by geography, then warming-induced shallowing or deepening of distributions may be predictable from species’ current depth syndromes and tolerance breadths. In contrast, predicting latitudinal changes in range size may require explicit representation of regional circulation, boundary currents, biogeographic breaks and habitat discontinuities, particularly for coastal and demersal taxa whose ranges are truncated by continental geometry. The partial coupling between depth and latitudinal ranges implies that species invading new latitudes may or may not expand vertically in parallel, depending on their traits.

Our study also underscores the importance of incorporating the vertical dimension into global macroecology. Much of the classical biogeographic literature treats marine species as effectively two-dimensional, described solely by their surface ranges. Yet our results show that vertical range is both systematically structured and tightly linked to depth, and that ignoring it can obscure strong Rapoport-like patterns and the ecological differentiation of taxa. The hypervolume of a marine species is inherently three-dimensional; range size in one dimension cannot be fully understood without reference to the others.

Our conclusions should be interpreted in light of several limitations. First, although our data set is large in absolute terms, it covers only ∼22,000 species and is biased toward vertebrates, especially fishes, so inferences about vertical and latitudinal range structure in poorly known invertebrate groups remain tentative. Second, the deep ocean is chronically under-sampled: both occurrence data and trait information are sparse below the bathyal zone, which likely leads to conservative estimates of richness and may preferentially detect wide-ranging “usual suspects” while missing narrow-ranged deep endemics. Third, our two key range dimensions are themselves products of heterogeneous data streams: vertical ranges are compiled from the literature and are formally independent of the latitudinal ranges derived from AquaMaps 2.0 modelled distributions, yet both are subject to observational and methodological biases. Spatial smoothing and environmental extrapolation in species distribution models may overestimate latitudinal extents, whereas depth limits drawn from regionally uneven literature could either under- or overestimate vertical ranges depending on where sampling effort has been concentrated. Fourth, our binning decisions in latitude and depth, and the use of simple linear relationships, may miss non-linear or scale-dependent components of Rapoport patterns. Future analyses could integrate finer-scale oceanographic structure, phylogenetic information and explicit measures of environmental tolerance to disentangle evolution from ecological contributions to range expansion. Nonetheless, the consistent vertical signal and fragmented horizontal signal uncovered here strongly suggest that, in the marine realm, Rapoport’s rule does apply but primarily in another dimension than historically defined.

## Data Accessibility Statement

All data used for this research will be available on Researchgate.

## Annexes

**Figure S1.**
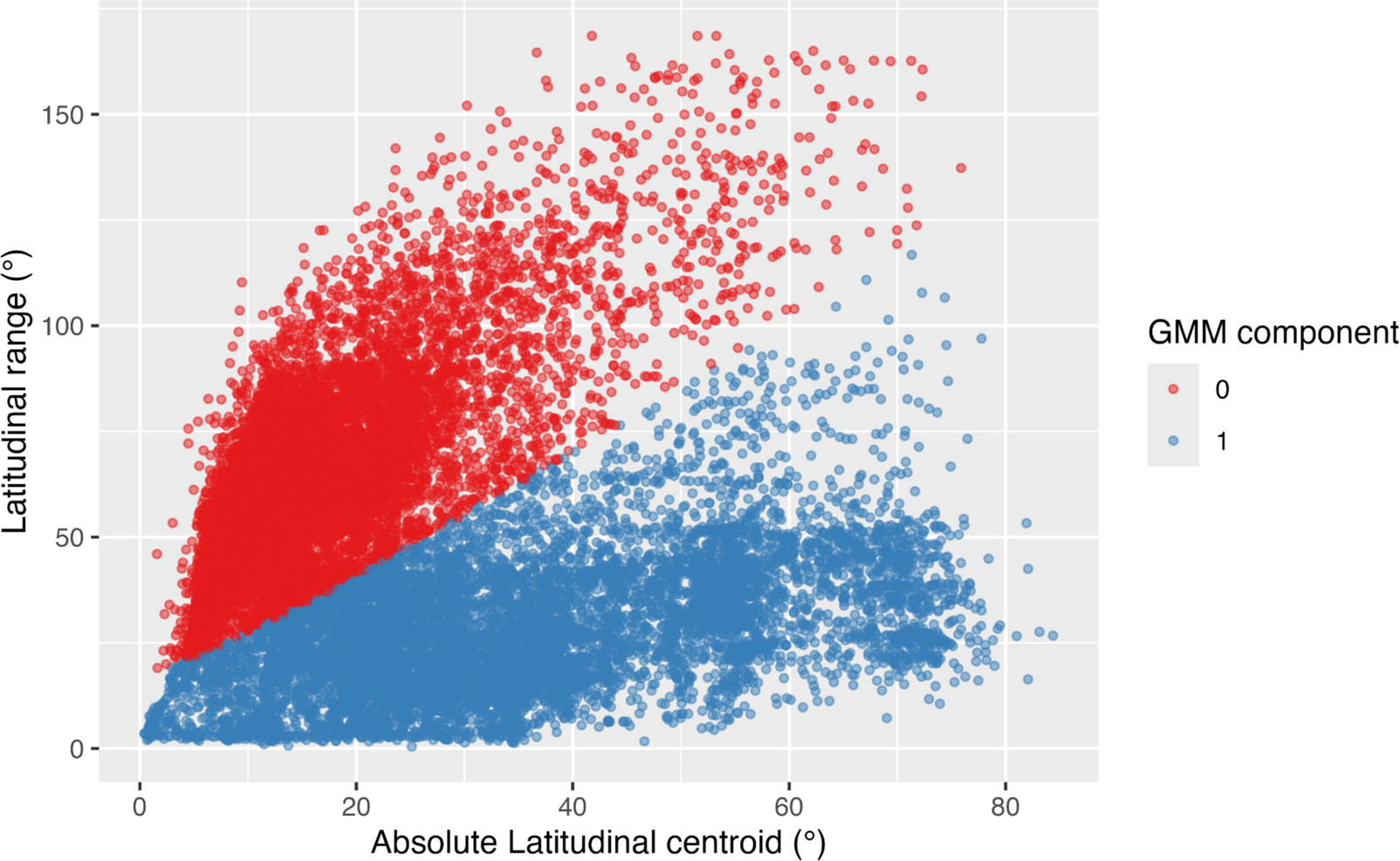
Mixture-model clustering of latitudinal ranges along the mid-latitude gradient. Scatter plot of latitudinal range size (°) as a function of mid-latitude (°) for all marine species in the dataset. Each point is a species, coloured by its posterior assignment to one of two components of a bivariate Gaussian mixture model fitted in (|mid latitude|, latitudinal range) space (component 0 in red, component 1 in blue).

## Notes

### Competing Interest Statement

The authors have declared no competing interest.

